# Sex Effect of E-liquid Constituents on Atherosclerosis and Gut Dysbiosis in ApoE^-/-^ Mice

**DOI:** 10.64898/2026.04.28.721512

**Authors:** Ann Marie Centner, Leila Khalili, Vladimir Ukhanov, Gwoncheol Park, Saurabh Kadyan, Hyun Seok Hwang, Ravinder Nagpal, Gloria Salazar

**Affiliations:** Department of Physiology, University of Tennessee Health Science Center, Memphis, TN 38112, USA; Department of Health, Nutrition and Food Sciences, Florida State University, Tallahassee, FL, USA

**Keywords:** e-cigarettes, menthol, atherosclerosis, inflammation, gut microbiome

## Abstract

**Rationale:** The role of sex in the effects of vaping and individual aerosolized e-liquid constituents on atherosclerosis, vascular aging, and gut microbiome remodeling remains poorly characterized.

**Objective:** To determine the contribution of e-cigarette aerosol components to vascular senescence, atherosclerosis, and gut microbiome dysbiosis in ApoE^-/-^ mice and vascular smooth muscle cell (VSMC) viability and senescence.

**Methods:** Male and female ApoE^-/-^ mice were exposed to e-liquid constituents (vehicle, vehicle plus nicotine, and vehicle plus nicotine plus menthol) for 48 minutes per day, 5 days per week for 16 weeks, with vascular pathology assessed in vivo. VSMCs isolated from aortas of wild-type and ApoE^-/-^ male and female mice were exposed to aerosolized e-liquids and evaluated for cellular senescence.

**Results:** Exposure to all tested e-liquid formulations, including vehicle, nicotine-containing, and menthol-containing aerosols, increased atherosclerosis in both male and female mice, with the most robust effects observed in the nicotine-containing formulation and in the descending aorta. Females exhibited greater sensitivity to e-liquid exposure, with increased plaque accumulation in both the aortic arch and descending aorta, while the addition of menthol was associated with reduced plaque burden compared with nicotine alone in both sexes. Novel findings show that e-liquid exposure also altered gut microbial composition in a sex- and exposure-dependent manner, with nicotine causing the greatest dysbiosis and menthol exerting modulatory, but not restorative, effects. Notably, Alloprevotella emerged as a key discriminating genus associated with reduced plaque burden, supporting a potential link between gut microbial remodeling, inflammatory regulation, and atherosclerosis.

**Conclusions:** These findings demonstrate that individual e-liquid aerosol components increase atherosclerosis and alter the gut microbiome in a sex-specific manner, with nicotine producing the most pronounced pro-atherogenic effects and the addition of menthol reducing these effects, without eliminating overall atherosclerotic risk.

## Introduction

Cardiovascular diseases (CVD) remain the leading cause of morbidity and mortality in the United States, with established sex differences^1^. Males more commonly develop coronary heart disease, whereas females have a higher incidence of stroke^2^. CVD risk increases with age, but lifestyle factors modify progression^1^. Tobacco cigarette use is the number one modifiable risk factor for CVD^3^ and exacerbates atherosclerosis and vascular senescence^4^. Nicotine acutely increases heart rate and vascular stress^5^, yet its direct contribution to atherosclerosis and cellular aging remains incompletely defined. With the rapid rise in nicotine-containing e-cigarette use— often perceived as less harmful than combustible cigarettes—clarifying their cardiovascular effects is increasingly important. Yet most studies have focused on pulmonary toxicity with limited investigation of vascular outcomes^6,7^.

Atherosclerosis is a chronic inflammatory condition involving endothelial cells (ECs), vascular smooth muscle cells (VSMCs), and macrophages. Initiation depends on lipid infiltration and oxidation within regions of endothelial dysfunction, leading to neointimal expansion.^8^ VSMCs undergo phenotypic switching, migrate into lesions, and can acquire a senescent phenotype that contributes to weakening of the fibrous cap and plaque destabilization, thereby accelerating disease progression^9,10^. Prior work implicates nicotine in vascular pathology through oxidative stress and inflammatory signaling; however, e-cigarette aerosols also contain vehicle and flavoring components that may independently affect vascular aging^4^.

Alterations in the gut microbiome have emerged as important contributors to CVD through effects on inflammation, microbial metabolites, and cardiometabolic risk^11^. Beyond microbial composition, intestinal barrier dysfunction and mucosal inflammation may also contribute to cardiovascular disease by increasing intestinal permeability and immune activation^12,13^. Despite growing interest in microbiome–cardiovascular interactions, the effects of e-cigarette aerosol exposure on gut microbial composition, intestinal barrier-associated pathways, and their potential relationship to cardiovascular outcomes remain incompletely explored.

Here, we investigated the sex-specific effects of e-cigarette aerosol constituents, including vehicle, nicotine, and menthol flavoring, on atherosclerosis, vascular senescence, and gut microbiome composition in atherosclerotic (ApoE^-/-^) mice. These *in vivo* studies were complemented by mechanistic *in vitro* experiments in WT and ApoE^-/-^ male and female VSMCs, to assess cellular senescence. Collectively, our findings demonstrate that all formulations exacerbate atherosclerosis, promote vascular senescence, and alter gut microbial networks, with nicotine driving the strongest effect on plaque burden and sex-specific differences observed across outcomes.

## Methods

### *In Vivo* Experiments

Male and female ApoE^-/-^ mice on a C57BL/6 background (3–4 months old; Jackson Laboratory) were randomized to control or aerosolized e-liquid exposure groups (vehicle, vehicle plus nicotine [24 mg/mL], or vehicle plus nicotine plus menthol) and exposed 48 minutes/day, 5 days/week for 4 months using a whole-body InExpose system (SCIREQ). Aerosols were generated from a 50:50 propylene glycol/vegetable glycerin vehicle using a temperature-controlled atomizer. A complete description of Materials and Methods, including primers for mRNA analysis (**Table S1**) and Major Resource Table (**Table S2**), is available in the Online Data Supplement.

## Results

### E-cigarette components regulate plaque content, cotinine, and serum lipids in a sex-specific manner in ApoE^-/-^ mice

Surprisingly, e-liquid exposure in male mice did not affect plaque content in the arch, while in females, arch plaque was increased by vehicle and nicotine (**Figure 1A and B**). In contrast, all e-liquid–exposed mice exhibited increased plaque area in the descending aorta (**Figure 1C**). Menthol attenuated nicotine-associated plaque accumulation in the aortic arch of females and in the descending aorta of both sexes. Aortic senescence-associated-β-galactosidase (SA-GLB1) increased with nicotine exposure in males and by all e-liquids in females (**Figure 1D**), indicating that menthol-associated plaque reduction occurred independently of senescence.

**Figure 1.**
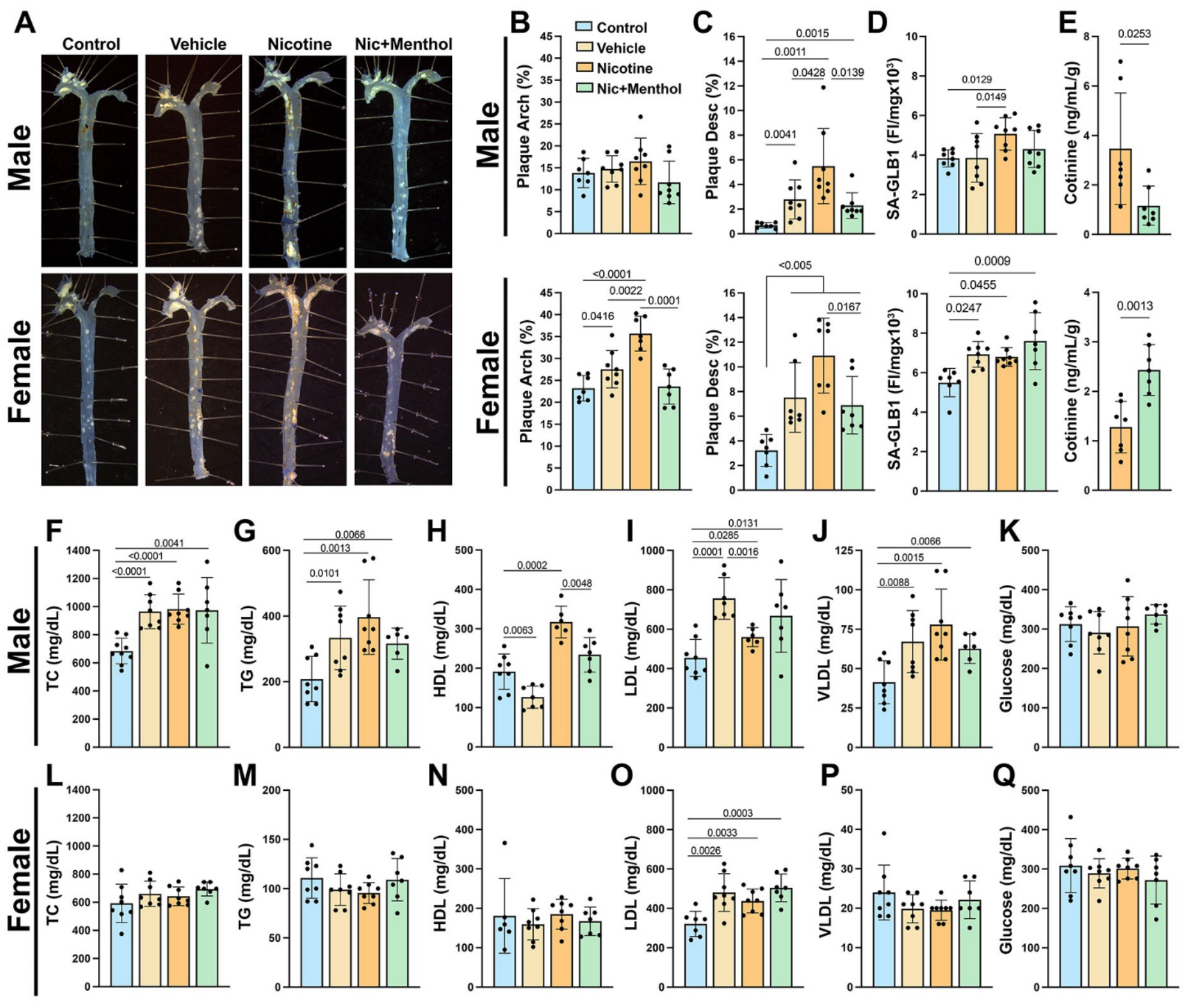
E-liquids increase atherosclerosis and LDL in male and female ApoE^-/-^ mice. Aortas from atherosclerotic mice were collected after treatment. Aortas were cleaned, then fixed and photographed after opening (A) for the analysis of the arch (B) and descending (C) plaque. Aortic senescence (SA-GLB1) was assessed using FDG (D), with a 12 h incubation. At week 16, plasma cotinine (E), lipids (F-J, L-P), including TC, TG, HDL, LDL, VLDL, and glucose (K, Q) were assessed. Significance of *P*<0.05 is indicated by the line between the 2 groups: n=6-9 per group for each sex. Significance was determined using a 2-way ANOVA with multiple comparisons.

To determine whether altered nicotine metabolism contributed to menthol-associated plaque reduction, serum cotinine was measured at study completion. Cotinine, a stable nicotine metabolite, is widely used to assess systemic exposure and allows cross-species comparisons. In humans, plasma cotinine levels in moderate smokers range from ∼100–300 ng/mL, while vaping produces a wide and variable range that can overlap with those observed in smokers^14-16^. Males exposed to nicotine alone had higher serum cotinine levels (61.1 ± 30 ng/ml) than those exposed to nicotine plus menthol (36.9 ± 26.4 ng/ml; p=0.15), but the difference did not reach significance. In females, cotinine was significantly lower in the nicotine (37 ±19.7 ng/ml) compared with the menthol group (59.4 ± 14 ng/ml, n=7, p=0.026). These differences were significant in both sexes after adjustment for body weight (**Figure 1E**). This finding is consistent with prior reports of elevated cotinine in female C57BL/6 mice exposed to mentholated versus non-mentholated cigarettes^17^.

Next, serum lipid, glucose, and liver enzyme concentrations were assessed. In males (**Figures 1F-J**), total cholesterol (TC), triglycerides (TG), low-density lipoprotein (LDL), and very-low-density lipoprotein (VLDL) were increased across all e-liquid groups, while high-density lipoprotein (HDL) was reduced with vehicle, increased with nicotine, and unchanged with menthol. In females, TC, TG, HDL, and VLDL were unaltered across treatments (**Figures 1L-P**). Glucose was unchanged between groups for both sexes (**Figures 1K, Q**). Overall, LDL was the only lipid significantly increased by all e-liquids in both sexes.

Sex-based comparisons (**Table S3**) revealed greater total plaque burden and aortic senescence in females across all exposure groups. Cotinine (adjusted by body weight) was higher in males than in females in the nicotine group, and higher in females than in males in the menthol group. Most lipids were significantly higher in males than in females across all groups, except for TC, HDL, LDL, and VLDL in controls and HDL in the vehicle group, which did not differ between sexes. An upward trend in TG in controls and in LDL in the menthol group was observed. No sex differences in glucose were observed. Collectively, sex-specific differences were evident in plaque burden, senescence, nicotine metabolism, and lipid profiles, with increased plaque in females associated with elevated senescence and altered nicotine metabolism rather than an adverse lipid profile.

Given the prominent sex difference in senescence, we exposed aortic vascular smooth muscle cells (VSMCs) from male and female C57Bl/6 wild-type (WT) and ApoE^-/-^ mice to e-cigarette extracts (vehicle, nicotine, and nicotine plus menthol). In both male cells, all extracts increased SA-GLB1 activity (**Figure 2A and B**). Female WT cells showed no response to vehicle and heightened activity with the other two formulations. Female ApoE^-/-^ cells responded differently, with increased activity across formulations. Notably, in female ApoE^-/-^ cells, senescence mirrored *in vivo* plaque findings, with the highest activity in the nicotine group. Across genotypes, female VSMCs exhibited higher SA-GLB1 activity than males, except in WT vehicle-treated cells (**Table S4**).

**Figure 2.**
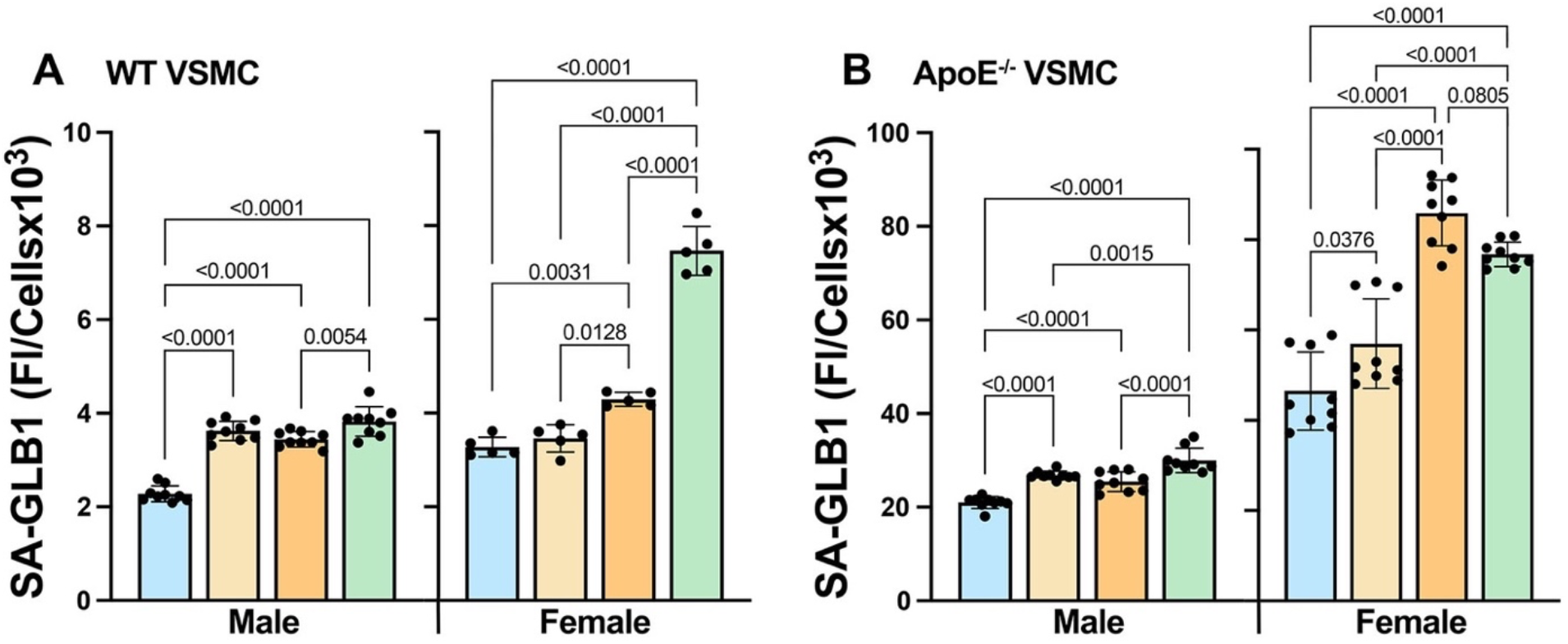
E-liquids increase senescence in male and female VSMCs. Male and female VSMCs (WT and ApoE^-/-^) were treated with 25% vehicle, nicotine, and nicotine plus menthol extracts for 3 d, and SA-GLB1 activity was assessed (A-B). Males n=9 and females n=5-9. Data is presented as mean ± SD with *P*-values <0.10 noted.

### Differences in atherosclerosis are independent of body composition and food intake after e-liquid exposure

To determine whether sex differences in plaque burden were driven by energy intake or body composition, food and water intake, body weight, fat mass, and lean mass were assessed. In males, body weight trajectories were similar between control and vehicle groups, while nicotine- and menthol-exposed males gained more weight than controls at the end of the intervention (week 16) compared with baseline (week 1) (**Figure S1A**).

Fat mass increased in all male groups at week 16 compared to week 1 (**Figure S1B**), while lean and water masses decreased for all male groups at week 16 compared to week 1 (**Figures S1C and D**), with the most significant decreases observed in nicotine-and menthol-exposed males. Importantly, these changes were not associated with increased food or water intake (**Figures S2A and C**), even after normalization to body weight (**Figures S2G and I**).

Female mice had comparable body weights across groups; despite weight gain in all groups at week 16 compared with week 1 (**Figure S1E**). No significant differences were seen for fat, lean, or water masses (**Figures S1F-H**). Across most weeks, food intake in all e-liquid– exposed female groups was higher or trended higher than control **(Figure S2B)**, with the nicotine-exposed group eating the most food. Water intake did not differ between female groups except at week 16 **(Figure S2D)**. These patterns persisted after body weight normalization **(Figures S2H and J)**, suggesting that increased food intake may explain weight gain in females but not in males.

Sex-based comparisons (**Table S3**) showed higher baseline and final body weight, lean mass, and water mass in males, while fat mass was higher in females. However, no differences in body composition between sexes were observed within the treatment groups by week 16. Heart rate was unchanged by treatment or sex (**Figure S3, Table S3**). Taken together, these results suggest that variations in plaque burden between sexes cannot be attributed to differences in food consumption, body weight, or body composition.

### E-liquids promote sex-specific inflammatory responses

Markers of systemic inflammation and organ weights were assessed to evaluate exposure-related physiological stress. In males, alanine transaminase (ALT) was reduced by vehicle and increased in the nicotine and menthol groups relative to controls, while aspartate transaminase (AST) was unchanged (**Figures 3A and B**). In females, ALT was unaffected, while AST increased only with menthol exposure. Liver weight increased with all e-liquids in both sexes, except in menthol-exposed males (**Figure 3C)**. For the spleen, no differences were seen in males, while in females, its weight was elevated in the nicotine and menthol groups, reaching significance only for nicotine (**Figure 3D**). The small intestine length was shorter in males exposed to nicotine, with no differences in females (**Figure 3E**). The same tendency (e.g., reduction in male nicotine group) was seen for cecum weight (**Figure 3F**). Overall, Inflammatory markers and organ measures exhibited sex- and exposure-specific effects.

**Figure 3.**
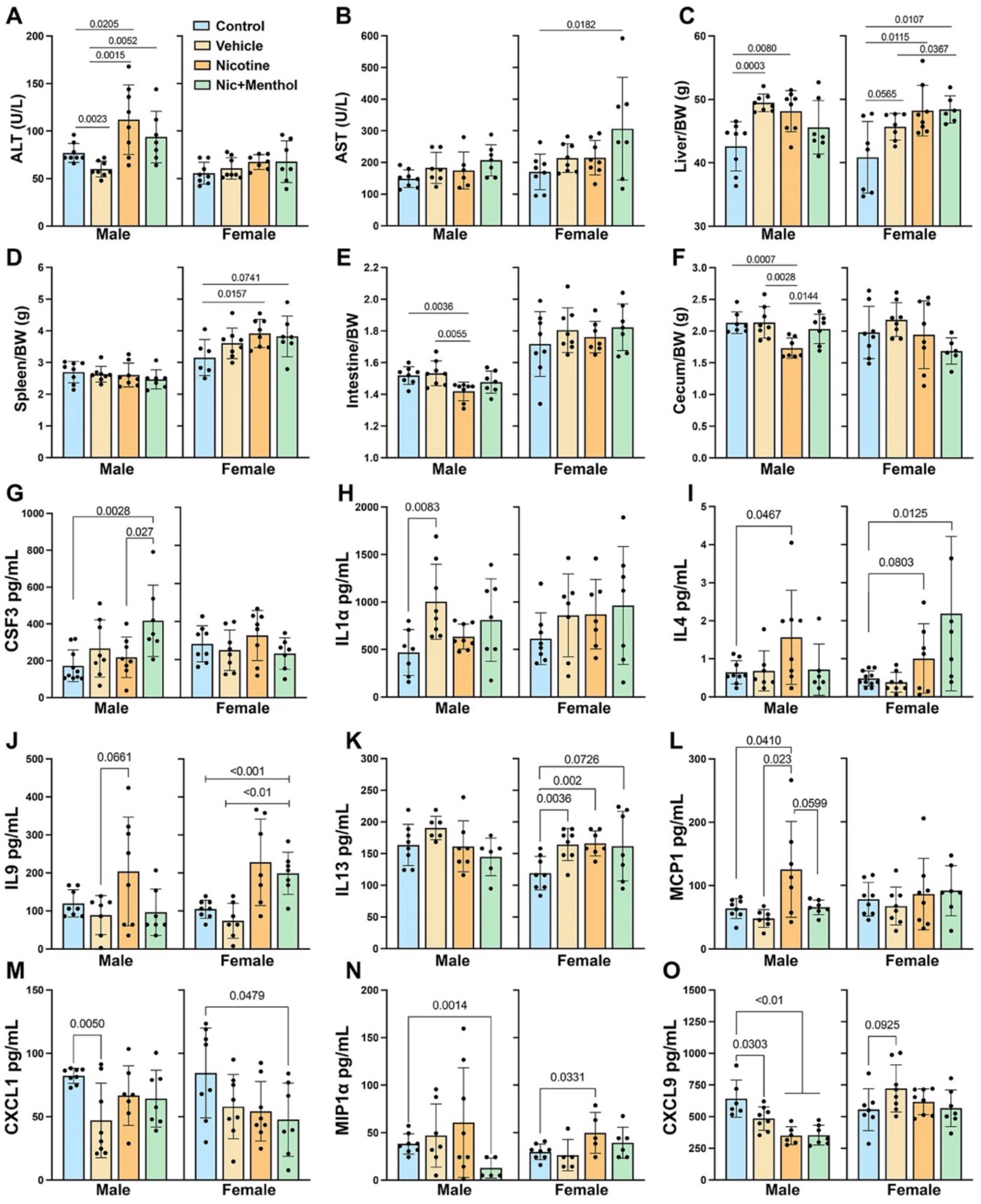
Sex effect of e-liquids in inflammation indices in ApoE^-/-^ mice. After 16 weeks of e-liquid exposure, serum liver enzymes (AST and ALT), and the liver, spleen, small intestine, and cecum were weighed and measured (A-F) and were divided by each animal’s body weight (BW). Serum inflammatory cytokines and chemokines were assessed using Luminex technology (G-O). Data is represented as mean ± SD (n=5-9) per group for each sex. Significance was determined using a 2-way ANOVA with multiple comparisons.

To assess systemic inflammation, serum was analyzed for expression of key cytokines and chemokines. Colony-stimulating factor 3 (CSF3), also known as granulocyte-stimulating factor (G-CSF), is a member of the interleukin-6 superfamily and controls granulocyte cytokine production, differentiation, and function^18^. It has been shown to be atheroprotective^19^ and was significantly higher in males but not in females in the menthol group (**Figure 3G**).

Interleukin 1 alpha (IL1α) is a potent inflammatory cytokine and senescence instigator upregulated by a milieu of oxidative and lipid overload stressors^20^. While IL1α trended higher across all e-liquids in both sexes, it reached significance only in vehicle-exposed males (**Figure 3H**). IL4 is upregulated during aging and cardiac fibrosis^21^. Here, IL4 was increased in both sexes exposed to nicotine and in females in the menthol group (**Figure 3I**). IL9 promotes atherosclerosis by increasing vascular cell adhesion molecule 1 (VCAM-1)-associated lesion inflammation^22^. In males, IL9 trended higher with nicotine, while in females, IL9 was elevated in the nicotine and menthol groups (**Figure 3J**). In a murine model, IL13 protects against atherosclerosis by altering macrophage type and plaque morphology^23^. While IL13 levels were stable in males, all three e-liquids elevated IL13 in females (**Figure 3K**). The cytokine monocyte chemoattractant protein-1 (MCP-1)/chemokine ligand 2 (CCL2) is secreted by VSMCs and ECs, recruiting monocytes to the lesion^24^. In males, MCP1 was elevated by nicotine and reduced with menthol, and its expression was unaltered in females (**Figure 3L**). Chemokine ligand 1 (CXCL1) is released by ECs and promotes atherosclerosis^25^. Interestingly, CXCL1 was reduced by vehicle in males and by menthol in females (**Figure 3M)**. The pro-inflammatory macrophage inflammatory protein-1 alpha (MIP1α)/chemokine ligand 3 (CCL3) is secreted by macrophages^26^, and it was reduced by menthol in males and increased by nicotine in females (**Figure 3N**). CXCL9 enhances monocyte adhesion to the endothelium during plaque formation^27^. CXCL9 was lower in all male groups exposed to e-liquids, while in females, it trended higher in the vehicle group (**Figure 3O**).

For sex comparisons within treatment groups (**Table S3**), ALT was higher in males in the control and nicotine groups. AST and cecum weights were unchanged between the sexes. Liver weight was higher in males in the vehicle group. In contrast, spleen weight and intestinal length were higher in females for all e-liquid groups. No significant sex effects were observed for CSF3, IL1β, IL4, IL9, MPC1, and CXCL1. A strong trend toward higher IL13 levels was observed in male controls. Females showed elevated MIP1α in the menthol group and higher CXCL9 in all e-liquid groups compared with males. Together, these results suggest that male atheroprotection is multifactorial and sex specific.

### E-cigarettes promote sex-specific changes in the microbiome

Given the microbiome’s role in systemic inflammation and cardiovascular health, we investigated whether nicotine induces microbiome dysbiosis and whether menthol attenuates this effect. We analyzed microbiome composition by 16S rRNA metagenomics in the feces of mice exposed to e-liquids. Vehicle (Veh), nicotine (NV), and nicotine plus menthol flavor (FLV) exposure each induced significant alterations in gut microbial beta-diversity, as observed in combined male and female data and when analyzed by sex (**Figure 4A**). The vehicle alone induced a significant separation in overall community structure from the control and nicotine groups. The greatest overlap in beta-diversity was observed between nicotine (NV) and menthol-flavored (FLV) groups, indicating more similar community compositions between these exposures. Alpha-diversity analyses revealed limited treatment effects. Chao1 richness was increased only in menthol-exposed males compared with controls, with no differences observed in other male or female groups (**Figure 4B**). Shannon diversity was increased only in nicotine-exposed females, while no significant differences were detected in males or other female e-liquid groups.

**Figure 4.**
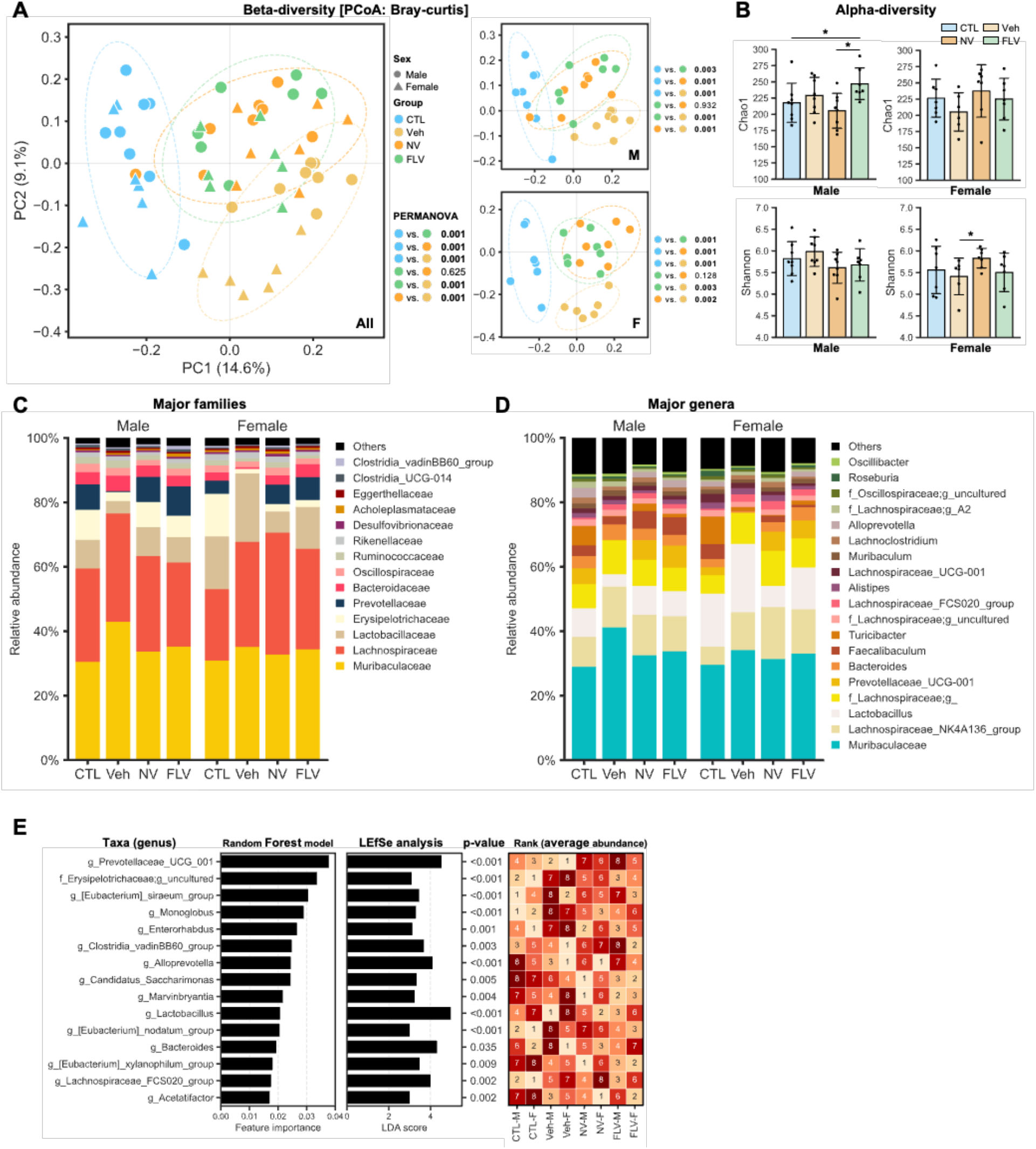
Sex-specific changes in gut microbial communities induced by E-liquids. At the completion of the experiment (4 months), fecal samples were collected from mice (n = 8 per group per sex) for microbiome analysis. (A) Beta-diversity and (B) alpha-diversity analyses demonstrate significant sex-dependent differences in response to Vehicle (Veh), nicotine, menthol flavor (FLV), and nicotine vehicle (NV) exposures compared with the control (CTL) group in males (M) and females (F). Alpha-diversity differences were assessed using the Kruskal-Wallis test, whereas beta-diversity significance was determined by permutational multivariate analysis of variance (PERMANOVA). Microbiome composition is shown at the (C) family and (D) genus levels. (E) Top 15 discriminative microbial features identified using a Random Forest model. The middle panel displays linear discriminatory analysis effect size (LEfSe) analysis comparing all four groups, and the right panel presents a heatmap showing abundance rankings based on the average relative abundance within each group. *p < 0.05 indicates statistical significance.

At the family level (**Figure 4C**), baseline sex differences were observed in controls, with females exhibiting lower Lachnospiraceae and Oscillospiraceae, and higher Lactobacilliaceae and Erysipelotrichaceae, than males. Vehicle exposure showed a stronger upregulation of Muribaculaceae in males than females and an upregulation in Lactobacilliaceae in females, with a reduction in males relative to sex-matched controls. Erysipelotrichaceae and Prevotellaceae showed a robust reduction in both sexes following vehicle exposure. Nicotine exposure showed stronger effects in females than in males. It increased Lachnospiraceae, reducing Erysipelotrichaceae and Lactobacilliaceae in females but not in males. Menthol exposure showed a profile more like that of controls in males. In contrast, in females, the profile was more like the nicotine group.

At the genus level (**Figure 4D**), in males, vehicle treatment induced the strongest effect on bacterial abundance, upregulating several bacteria, including *g_Muribaculaceae, g_Lachnospiraceae_FCS020_group, and f_Erysipelotrichaceae;g_uncultured* and reducing *g_Turicibacter* and *g*_*Provatellaceae*_ *UCG_001* compared with controls (**Figure S4A**). Compared with the vehicle, nicotine increased *g_Provatellaceae*_ *UCG_001, g_Lactobacillus* and *g_Alloprevotella*, reducing several taxa, including *f_Erysipelotrichaceae;g_uncultured*, and *g_Akkermansia* (**Figure S4B**). Compared with Nicotine, menthol upregulated *g_Anaeroplasma, g_Clostridia_vadinBB60_group*, and *f*_*Oscillospiraceae;g_uncultured* (**Figure S4C**). Similar changes were observed in females, with fewer bacteria affected by the vehicle than in the control (**Figure S4D**). Compared with the vehicle, nicotine upregulated more taxa in females than in males (**Figure S4E**), including *g*_*Provatellaceae*_ *UCG_001, g_Clostridia_vadinBB60_group, g_[Eubacterium]_siraeum_group*, and *g_Ruminococcus*. Unlike males, no bacteria were upregulated by menthol compared with nicotine in females (**Figure S4F**).

A machine learning approach using a random forest model was employed to identify taxa contributing to differences among groups, and the results were supported by linear discriminant analysis **(Figure 4E)**. Among the 15 taxa with the highest feature importance for classification, *Prevotellaceae_UCG_001* emerged as the most important taxon, showing higher abundance in the Control and Vehicle groups compared with the Nicotine- and menthol-treated groups and a tendency toward higher abundance in females. This was followed by an uncultured genus within the Erysipelotrichaceae family. In addition, *Candidatus_Saccharimonas* was identified as another important feature, displaying relatively higher abundance in the nicotine-treated group.

### E-liquids induce sex-specific effects on intestinal inflammation and intestinal permeability

Because of the significant changes in microbiome composition induced by e-liquids, we measured inflammatory markers known to be upregulated in intestinal inflammatory conditions. Compared with controls, the vehicle increased IL1β and TNFα in males (**Figure 5A and F**) and IFNγ and MCP1 in females (**Figure 5G and H**). In males, nicotine upregulated IL1β and TNFα (**Figures 5A and F**), with an upward trend in IL8 and IL10 (**Figures 5C and D**). In contrast, in females, nicotine increased IL6, IFNγ, and MCP1 (**Figures 5B, G, and H**). Menthol reduced nicotine effects in males for IL1β, IL8, IL10, IL17, IFNγ, and MCP1, with a downward trend for TNFα (**Figures 5A, C-H**). In females, menthol increased IL17A, IFNγ, and MCP1 (**Figures 5E, G, and H**). IL6 was reduced compared with nicotine, but it did not reach significance (**Figure 5B**). No treatment effects were seen for IL6 in males and IL1β, IL8, IL10, and TNFα in females, although TNFα trended higher with menthol (**Figures 5B, A, C, D, and F**).

**Figure 5.**
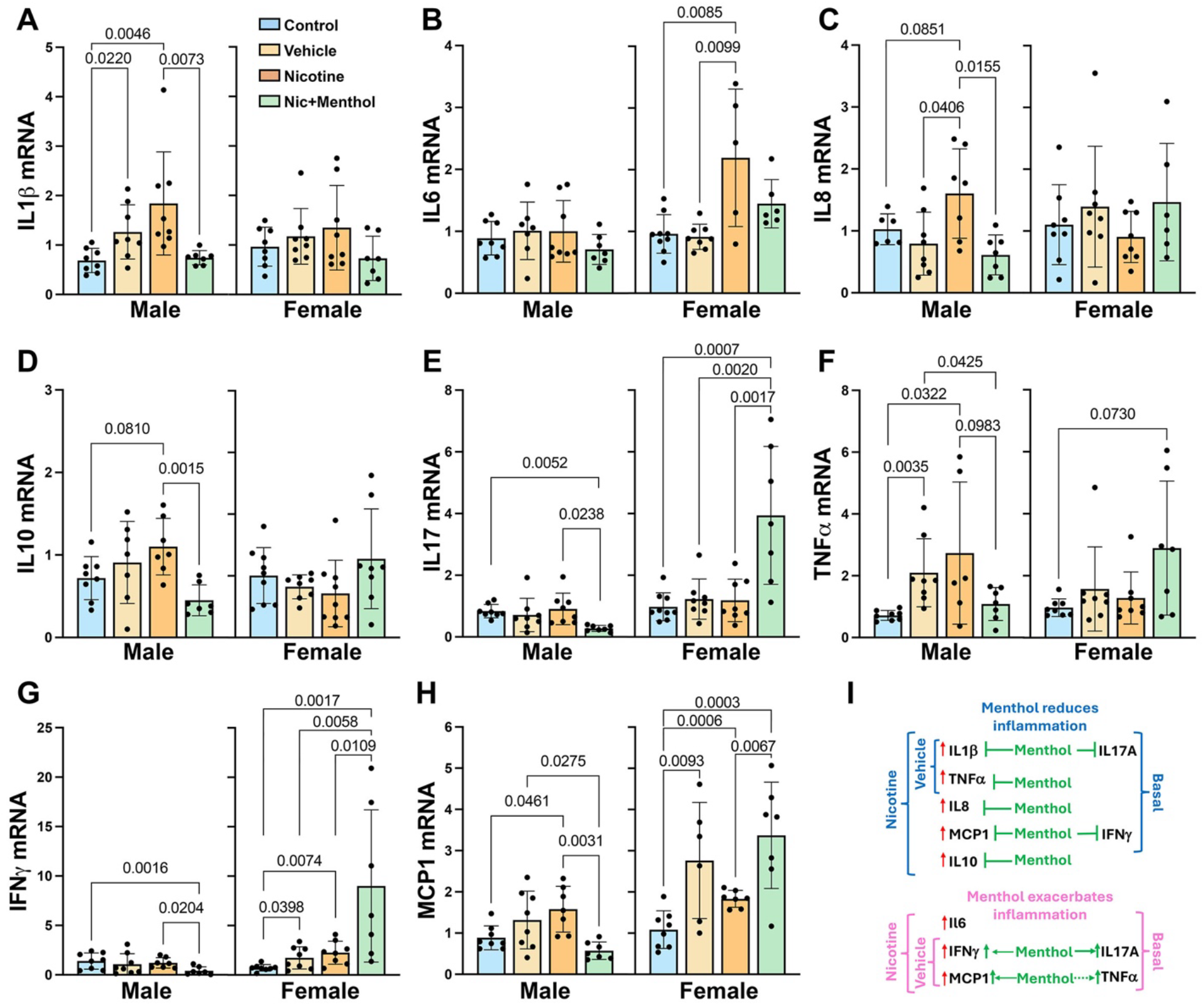
Sex effects in inflammatory response to E-liquids. After 4 months of e-liquid exposure, the ileum from male and female atherosclerotic mice was collected and processed for mRNA expression of inflammatory markers (**A-H**). Data is represented as mean ± SD (n=5-9) per group for each sex. Significance was determined using a 2-way ANOVA with multiple comparisons. In males (blue), menthol appeared to reduce the inflammatory burden, whereas in females (pink), it exacerbated inflammation (**I**). Arrows denote increased effects with -l designates blunted effects. Dashed line designates trends, while undashed lines were indices that reached significance.

The data summarized in Figure 5I indicates that nicotine promotes intestinal inflammation in both sexes, which was reduced by menthol in males and was exacerbated by menthol in females.

For sex effects (**Table S5**), females showed an upward trend for MCP1 in the vehicle group and significantly higher IL6 levels in the nicotine group. Menthol elevated IL6, IL17A, IFN,γ and MCP1 in females compared with males. A trend toward higher levels was also seen for IL10.

The sex-divergent effects of menthol on inflammatory cytokines in the ileum suggest that menthol may have sex-specific effects on intestinal permeability. To this end, we measured mRNA expression of tight junction proteins and mucin genes. In males, nicotine and vehicle reduced zonula occludens 2 (ZO2) and junctional adhesion molecule 3 (JAM3), whose levels were restored to baseline levels by menthol (**Figures 6B and D**). Occludin (OCLN) expression was unaffected by the e-liquids, while ZO1, claudin 1 (CL1), and CL2 expression were only affected by menthol, which increased their levels compared with controls **(Figure 6A, C, E, F**). Regarding mucin (Muc) genes, nicotine upregulated Muc6, which was reduced by menthol (**Figure 6H**). No treatment effects were seen for Muc2 and Muc13 (**Figure 6G and I**).

**Figure 6.**
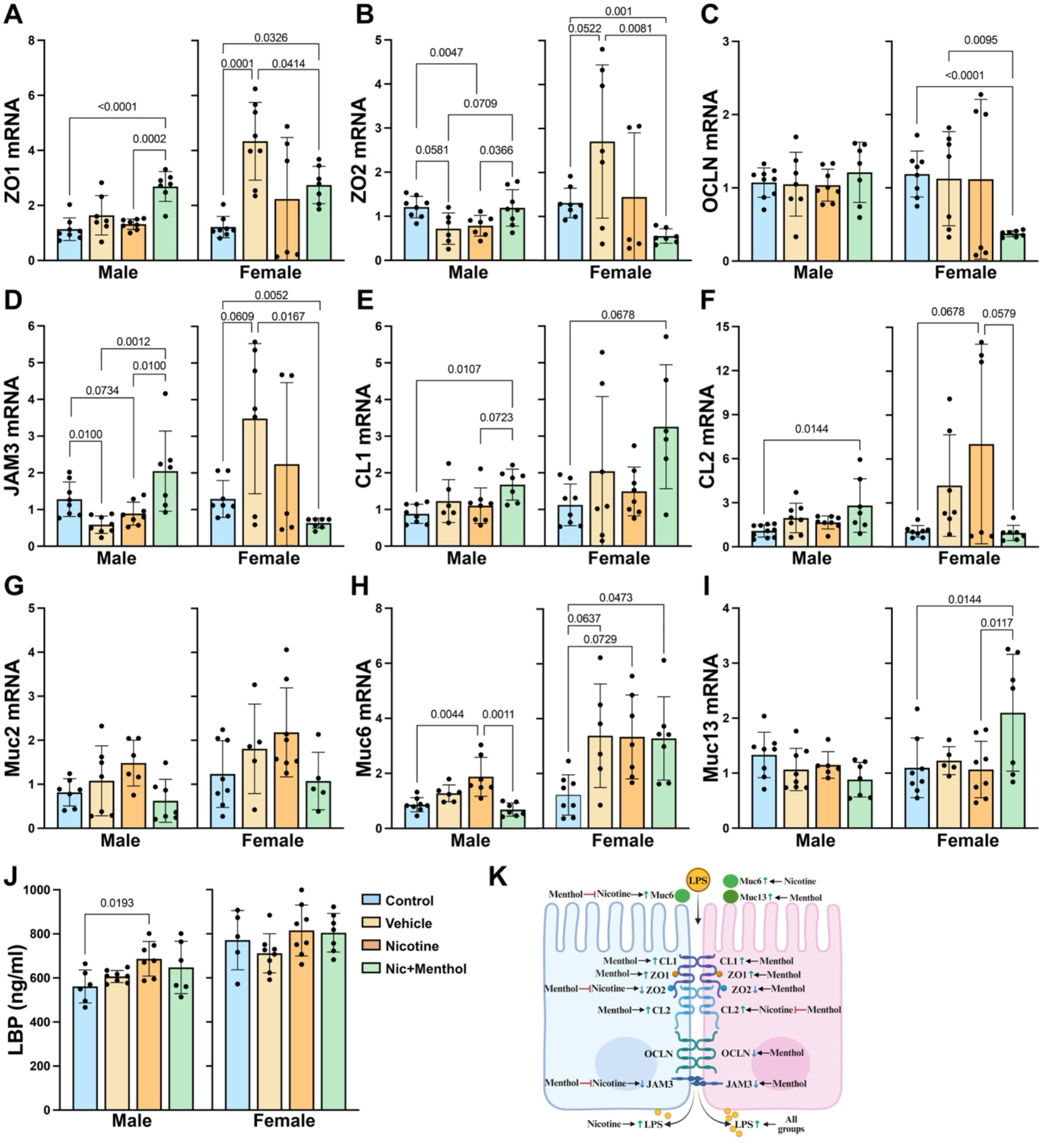
Sex effects of E-liquids on intestinal epithelial tight junction gene expression and endotoxemia. The ileum of male and female mice exposed for 4 months to e-liquids was collected and processed for mRNA expression of tight junction proteins (A-F) and mucin genes (G-I). Endotoxemia was assessed by measuring serum LBP. The model depicted in K summarizes the regulation of intestinal barrier components under different exposures. Data are presented as mean ± SD, with sample sizes ranging from n = 5-9 per group.

In females, ZO1, ZO2, and JAM3, CL1 and CL2 were increased by vehicle, reaching significance only for ZO1 (**Figure 6A, B, D, E, F**). Nicotine exposure led to an upward trend for CL2, while menthol exposure reduced ZO2, OCLN, and JAM3 compared with control and vehicle (**Figure 6F, B-D**). ZO1 was reduced by menthol compared with the vehicle but was elevated compared with the control. Like males, CL1 was elevated by menthol in females (**Figure 6E**). CL2 was reduced by menthol compared with nicotine, reaching control levels (**Figure 6F**). While Muc2 was not altered by treatment, Muc6 was elevated by all e-liquids compared with control, reaching significance for menthol (**Figures 6G and H**). Muc13 was elevated by menthol compared with control and nicotine (**Figure 6I)**.

Playing a key role in the innate immune response, lipopolysaccharide (LPS) binding protein (LBP) mediates the inflammatory response to LPS^28^ and serves as a marker of intestinal permeability. LBP was elevated in males exposed to nicotine, while no treatment effects were seen in females **(Figure 6J**). Data summarized in **Figure 6K** support a model in which nicotine exposure increases epithelial permeability in males, potentially by reducing ZO2 and JAM3, accompanied by elevated LBP in circulation. Menthol exposure in males ameliorated these effects by increasing ZO1 and JAM3. The increase in the pore-forming claudin CL2 in females with vehicle and nicotine suggests that LPS and/or other inflammatory molecules may leak into circulation, as CL2 promotes inflammation in a mouse model of colitis^29^. In females, menthol significantly reduced ZO2, OCLN, and JAM3, suggesting exacerbated intestinal permeability.

For sex differences (**Table S5**), compared with males, females had elevated ZO1, ZO2, JAM3, and Muc6 in the vehicle group, higher Muc6 in the nicotine group, and lower ZO2, OCLN, JAM3, and CL2 in the menthol group. Higher levels of Muc6 and Muc13 were also observed in females in the menthol group. No sex differences were noted at basal. Females had elevated LBP in all female groups,

We next evaluated the effects of e-liquid formulations on morphological changes (**Figure 7A**), as inflammatory conditions reduce villus height while increasing villus thickness and crypt depth^30^ (**Figure 7B**). Villi height was highest following vehicle exposure, was reduced by nicotine, and was restored to control levels by menthol in both sexes (**Figure 7C**). The villi thickness was elevated by nicotine and menthol in males and only by menthol in females (**Figure 7D**), while the crypt depth was elevated by all e-liquids in both sexes (**Figure 7E**). Additionally, goblet cell number was reduced by all e-liquid exposures in males but not females (**Figure 8A-C**).

**Figure 7.**
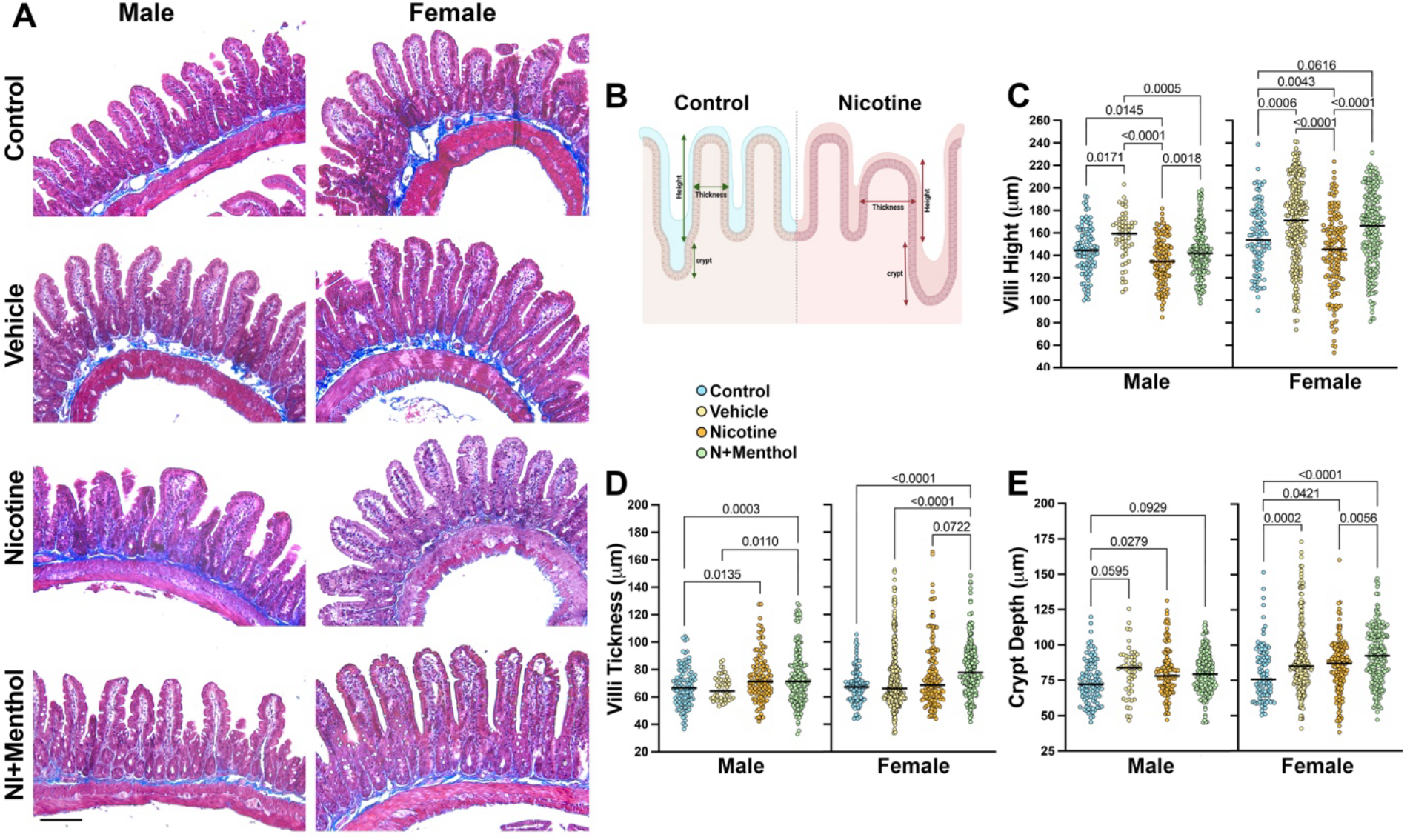
Sex effects of E-cigarette components on intestinal architecture. After 4 months of E-liquid exposure, the ileum from male and female mice was collected, fixed, and stained with H&E to assess villi morphology. Representative images of E-liquid-exposed and control mice (A). Model representing the change in villi structure exposed to (**A-E**). Data is represented as mean ± SD (n=5-9) per group for each sex. Significance was determined using a 2-way ANOVA with multiple comparisons.

**Figure 8.**
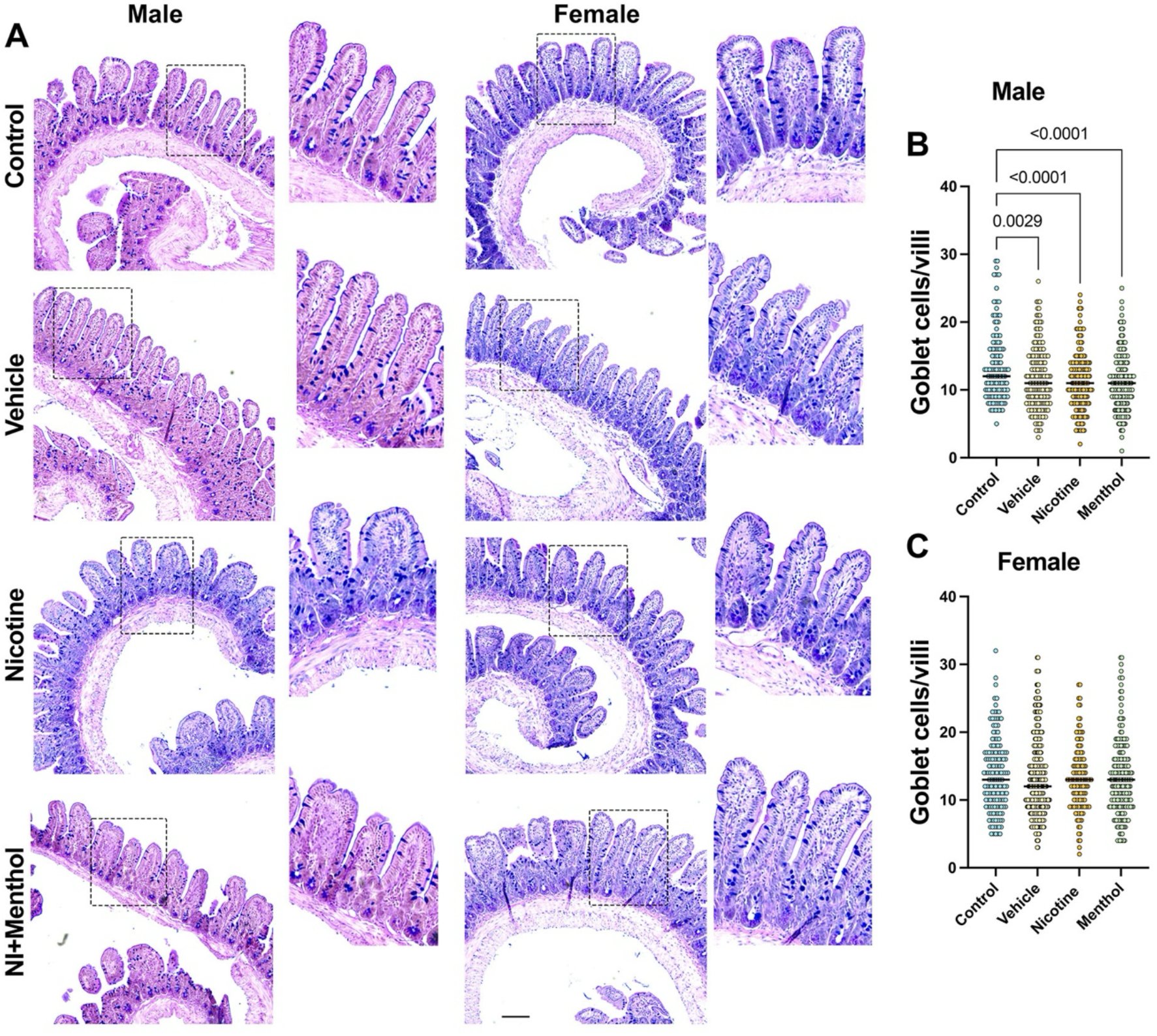
Effects of E-liquids exposure on ileal goblet cell abundance assessed by Alcian blue staining. Representative histological images of ileal sections from male and female mice from experimental groups (Control, Vehicle, Nicotine, and Nicotine + Menthol). Alcian blue staining highlights mucin-producing goblet cells (blue) within the intestinal epithelium. Insets show higher magnification views of villi. Quantification of goblet cells per villus in males (B) and females (C).

For sex effects (**Table S5**), villus height and crypt depth were higher in females than in males in all groups, while villi thickness was higher in females in the vehicle and menthol groups. Goblet cell numbers were higher in all e-liquid-exposed females.

### Associations among gut microbial taxa, inflammation, and atherosclerosis

To investigate the correlation between bacterial abundance and physiological and biochemical parameters, correlation analyses were performed (**Figure 9**). Notably, the genus *Enterorhabdus* and its family-level, Eggerthellaceae, exhibited multiple significant associations across a range of parameters. Specifically, *Enterorhabdus* showed positive correlations with serum IL1α and CXCL9, a chemokine induced exclusively by IFNγ, as well as with ileal IFNγ levels. In addition, it was positively associated with intestinal length and cecum weight. In contrast, *Alloprevotella* displayed an opposite pattern, showing negative correlations with ileal IL6 and with both aortic arch and descending atherosclerotic plaque burden. It was also negatively correlated with spleen weight, while showing positive associations with HDL levels and ALT. Notably, *Alloprevotella* abundance was reduced in both the vehicle and nicotine-treated groups but restored in the menthol-treated group (**Figure 4, S4**), closely mirroring the observed group differences in plaque burden (**Figure 1B-C**). Furthermore, differences in bacterial abundance between groups were more pronounced in females, consistent with the observed sex-specific differences in plaque formation. Beyond these taxa, the *Clostridia_vadinBB60_group* showed a negative correlation with aortic arch plaque, whereas the *Lachnospiraceae_FCS020_group* demonstrated a positive correlation with descending plaque burden.

**Figure 9.**
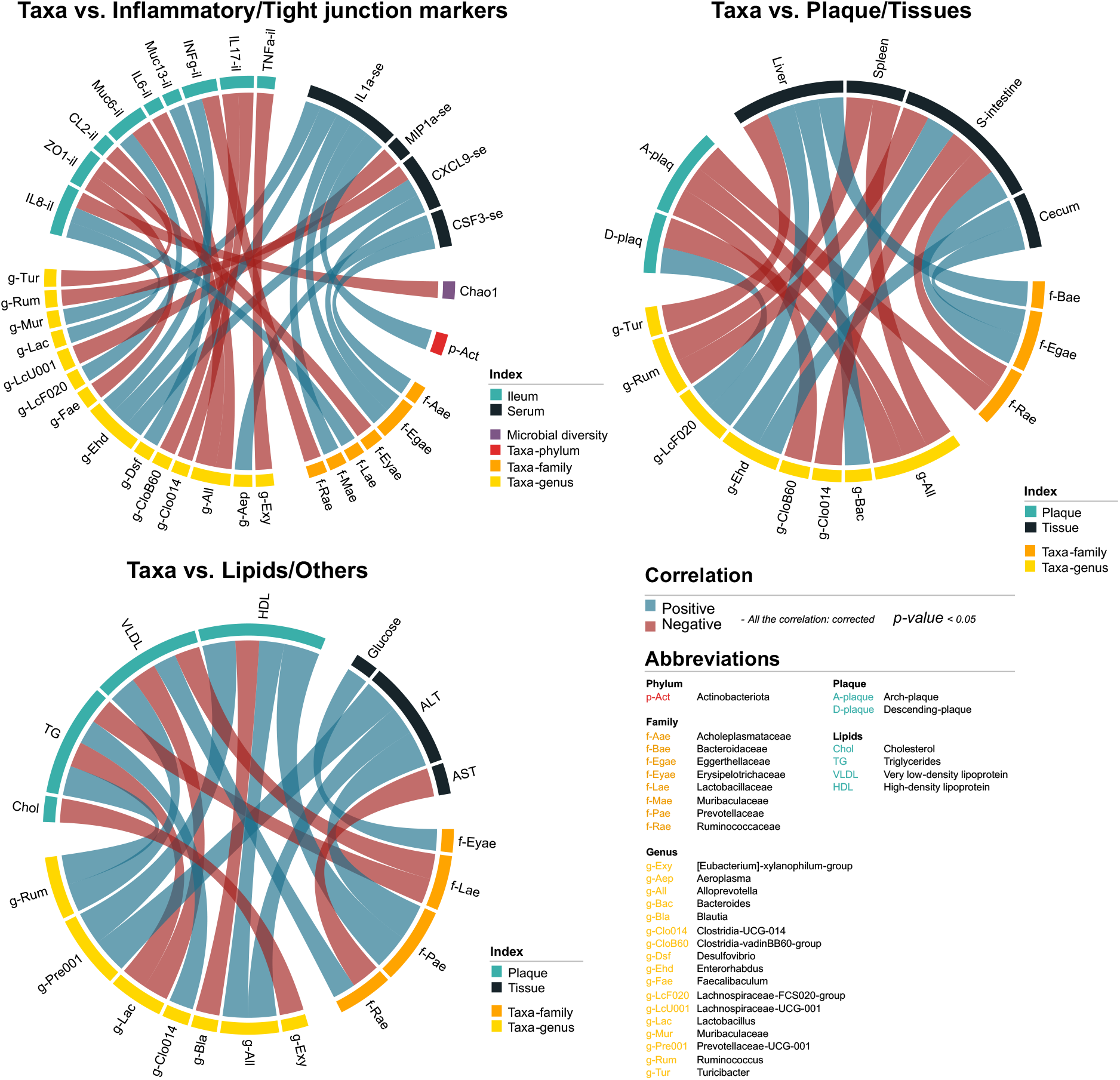
Correlation networks linking gut microbiota, inflammation, lipids, and atherosclerotic plaque burden. Spearman’s correlation analysis showing associations between microbial taxa and inflammatory/tight junction markers (upper left), plaques and tissue length/weight parameters (upper right), and lipid profiles and additional physiological parameters (lower right). Only statistically significant correlations (adjusted p < 0.05) are displayed. The width of each connecting line corresponds to the absolute value of the correlation coefficient. Blue lines indicate positive correlations, whereas red lines indicate negative correlations

Overall, e-liquid exposures disrupted intestinal inflammation, barrier-associated gene expression, and ileal morphology in a sex-dependent manner, with nicotine promoting intestinal dysfunction and menthol exerting divergent effects by sex. Correlation analyses further linked microbial taxa, particularly *Alloprevotella*, with inflammatory markers and plaque burden, supporting a potential association between gut microbial remodeling and cardiovascular outcomes.

## Discussion

This study extends prior reports describing adverse cardiovascular effects of aerosolized e-liquids^31^ by identifying sex-specific and constituent-dependent mechanisms linking nicotine, menthol, inflammation, metabolism, senescence, and atherosclerosis. Using complementary *in vivo* and *in vitro* approaches across both sexes, this work addresses a critical gap in understanding how e-cigarette constituents differentially influence cardiovascular risk.

Plaque outcomes were strikingly sex dependent. Increased plaque accumulation in the aortic arch was found exclusively in females following e-liquids, while descending plaque was observed for both sexes. These differences underscore the need for thorough vascular assessment in cardiovascular risk evaluation. Studies that focus solely on the aortic root potentially overlook region-specific effects. For example, a study that exposed ApoE^-/-^ mice to 12 weeks of 2.4% nicotine aerosol only analyzed a cross-section of the aortic root for atherosclerotic burden, in which there was a significant increase with treatment^40^. Vascular senescence patterns partially mirrored plaque distribution, with increases corresponding to descending plaque accumulation. Given evidence linking senescence to plaque instability^41^, these associations suggest potential mechanistic overlap.

To our knowledge, male and female WT and ApoE^-/-^ cell lines have not previously been directly compared in this context, highlighting the novelty of these findings. In males, aortic senescence increased only with nicotine exposure *in vivo*, whereas male VSMCs exhibited increased senescence across all e-liquid constituents *in vitro*, highlighting altered apoptotic resistance^67^ and the importance of whole-organism metabolism. In females, WT cells showed the highest senescence in response to menthol. In contrast, menthol exposure in ApoE^-/-^ cells resulted in a slight attenuation of senescence, highlighting genotype-specific effects and mirroring *in vivo* findings. The higher baseline senescence in female cells remains unexplained but is consistent with the more pro-atherogenic *in vivo* phenotype.

A central finding was sex-specific divergence in energy handling following e-liquid exposure, with increased intake and weight gain observed in females but not males. Nicotine-exposed males gained body weight by the study endpoint despite no increase in intake. Additionally, observed weight gains did not correlate with plaque burden, indicating that body weight alone does not predict atherosclerotic risk. This contrasts with prior work showing nicotine-associated suppression of weight gain^32-34^ and highlights an inconsistent relationship between e-cigarette exposure and body weight^35^. Although the relationship between e-cigarette exposure and body weight remains unresolved, evidence from combustible cigarette studies links heavy smoking to increased adiposity and insulin resistance—both established contributors to atherosclerosis and cardiovascular risk^36-38^.

Consistent with altered metabolic handling, males, but not females, exhibited increased fat mass and reduced lean mass over time—patterns that were not aligned with plaque burden. In males, nicotine may alter metabolism by blunting lipogenesis and fat oxidation, potentially via ROS production and AMPK phosphorylation, as previously describe^35^. These mechanisms could be further explored in aortic tissue or in VSMCs.

Given nicotine’s stimulant properties, we anticipated but did not see elevations in heart rate among the nicotine-exposed mice. While vaping increases heart rate in humans^39^, prior murine studies have reported decreased heart rate following short-term heavy vaping exposure^32^, suggesting species- and exposure-dependent cardiovascular responses.

Findings from this study and previous reports suggest that menthol flavoring affects nicotine metabolism^17,42^. Here, menthol exposure decreased cotinine levels in males but increased levels in females. Elevated cotinine in females may reflect slower nicotine clearance or enhanced conversion of nicotine to cotinine. Supporting this interpretation, menthol has been shown to upregulate CYP2A6—an enzyme involved in nicotine metabolism—in plasma extracellular vesicles^43^.

Prior work using nicotine-containing drinking water with and without menthol reported increased consumption without increased cotinine in menthol-exposed males, but not females^44^, suggesting sex-specific differences in nicotine metabolism or handling. Consistent with altered nicotine handling, menthol-exposed females in our study exhibited reduced plaque burden in the aortic arch. Human studies further support a role for menthol in modulating nicotine metabolism, as mentholated cigarette use is associated with reduced conversion of nicotine to cotinine compared with non-mentholated cigarettes^45^. Collectively, these observations raise the possibility that cotinine may exert sex-specific cardiovascular effects. Further studies are warranted to determine whether menthol-mediated modulation of nicotine-metabolizing enzymes, including CYP2A6 and flavin-containing monooxygenases (FMOs), contributes to these effects.

Changes in circulating lipids indicate that e-liquids may contribute to descending plaque accumulation through sex-specific mechanisms. LDL increased across all exposure groups and may contribute to plaque build-up in both sexes, whereas elevations in TC, TG, and VLDL in males did not align with increased plaque.

Vehicle exposure alone induced a pro-atherogenic lipid profile and plaque accumulation, supporting a contribution of the vehicle to cardiovascular risk. Combustion of PG and VG in the e-liquid vehicle generates reactive carbonyl compounds, including acrolein^46,47^. Acrolein can elevate TC and TG, induce atherosclerosis in male ApoE^-/-^ mice^48^, and impair HDL function in reverse cholesterol transport^49^, suggesting that HDL quantity may not reflect HDL functionality. Thus, observed increases in HDL may reflect compensatory responses rather than protection, particularly given evidence that lipoprotein oxidation amplifies atherogenicity ^50,51^.

The pro-inflammatory nature of atherosclerosis, combined with nicotine “stress” exposure, led us to anticipate systemic inflammation changes^20,25,52^. In our study, more inflammatory markers were upregulated in females than in males in the nicotine and menthol groups (**Table S6**). In males, more inflammatory molecules were reduced in the menthol group than in females. A recent meta-analysis identified CSF3/G-CSF as atheroprotective^19^, and this cytokine was increased in menthol-exposed males. In parallel, menthol exposure in males was associated with reductions in pro-fibrotic IL4, pro-inflammatory IL9, and other pro-atherogenic mediators, including MIP1α and CXCL9—effects not observed in females. Notably, increased CSF3, alongside decreased MCP1 and MIP1α, correlated with reduced descending plaque accumulation in males exposed to menthol plus nicotine, suggesting that these inflammatory shifts may affect plaque burden. MCP1 levels correlated with plaque in males, as it was increased by nicotine and reduced by menthol. Collectively, this profile is consistent with reduced monocyte recruitment, attenuated macrophage differentiation, decreased extracellular matrix remodeling, and potentially diminished lipid infiltration and oxidation within lesions.

In contrast, menthol-exposed females, who had reduced aortic plaque burden, showed increased IL4 and IL9 and decreased CXCL1. Elevated IL4 may reflect a shift toward a more stable plaque phenotype. While some inflammatory changes overlapped with nicotine exposure, the reduction in CXCL1 and elevation in IL9 were unique to menthol-exposed females and associated with less aortic plaque than nicotine. Notably, the only cytokine elevated in females exposed to nicotine was MIP1α. Given that MIP1α was reduced in menthol-exposed males and is associated with increased risk of fatal cardiovascular events in humans^53^, this chemokine may represent a key mediator of sex-specific plaque progression. Lastly, IL13 levels were unaffected in males but increased in females across all e-liquid exposures, mirroring the greater plaque burden in females. Whether this interleukin is protective is unknown^23,54,55^.

All e-liquid exposures altered gut microbial composition; however, nicotine induced the most pronounced restructuring, as reflected by strong beta-diversity separation across sexes. The close overlap between nicotine and menthol groups indicates that menthol does not normalize nicotine-induced dysbiosis but instead modifies community structure within a similarly perturbed state. Limited, sex-specific changes in alpha diversity suggest that microbiome remodeling occurs primarily through compositional reorganization rather than uniform shifts in richness or evenness, with menthol increasing Chao1 richness only in males and nicotine increasing Shannon diversity only in females.

At the taxonomic level, nicotine produced broad, sex-dependent disruption across phylum, family, and genus levels, including depletion of multiple short-chain fatty acid (SCFA)–associated taxa, particularly in females. Menthol partially preserved select taxa in a sex-dependent manner, yet did not restore coordinated microbial networks, consistent with a modulatory rather than restorative effect. Vehicle exposure alone also altered microbiome composition, underscoring contributions from non-nicotine constituents. Collectively, these findings indicate sex- and exposure-dependent network-level disruption of the microbiome, which aligns with observed differences in atherosclerotic plaque burden.

Our findings extend prior studies linking tobacco-related exposures to microbiome dysbiosis. In female ApoE^-/-^ mice, cigarette smoke exposure increased Akkermansiaceae—an effect reversed by smoking cessation—while aerosol exposure increased Lactobacillaceae^56^. Increased Akkermansiaceae has also been reported in male mice exposed to cigarette smoke ^57^. Although Akkermansiaceae is more abundant in healthy individuals compared with patients with inflammatory bowel disease^58^ and obese humans^59^, cigarette smoke exposure in these models is nonetheless associated with adverse outcomes, including atherosclerosis, lung inflammation, and emphysema^60^. While smoking has been linked to microbiome dysbiosis^61^, vaping has not yet shown uniform effects, underscoring the need for exposure- and sex-specific evaluation.

The partial plaque reduction in menthol-exposed females, together with the identification of *Alloprevotella* as a key discriminating genus, represents a novel finding. *Alloprevotella*, a commensal gut bacterium in humans and mice, has been increasingly associated with protection against inflammatory disorders^62,63^ and higher abundance with reduced cardiovascular disease risk^64^. Previous studies have reported that lycopene, a dietary carotenoid with antioxidant and anti-inflammatory properties, significantly attenuated atherosclerosis^65^. This effect was accompanied by increased *Alloprevotella* abundance and a negative correlation between *Alloprevotella* and inflammatory markers, including IL6^65^, consistent with the associations observed in the present study. Similarly, treatment with glycoursodeoxycholic acid, a bile acid metabolite with reported anti-inflammatory and microbiome-modulating effects, increased *Alloprevotella* abundance, which was inversely correlated with atherosclerotic plaque area^66^. Despite limited direct mechanistic evidence, accumulating findings support a plausible association between *Alloprevotella*, inflammatory regulation, and plaque progression, warranting further investigation into its role in atherosclerotic plaque formation and cardiovascular disease progression.

As with serum inflammatory molecules, in the ileum, several markers were downregulated in the menthol group compared with nicotine in males, and more genes were upregulated by menthol in females (**Table S6**). MCP1 expression in the ileum also correlated with plaque burden in males, while in females, MCP1 was elevated in all e-liquid-exposed groups. Nicotine exposure may affect gut barrier-associated pathways, as reflected by increased intestinal inflammatory signaling (including IL1β and TNFα in males and IL6, IFNγ, and MCP1 in females), reduced ZO2 and JAM3 with elevated circulating LBP in males, and sex- and treatment-dependent alterations in mucin expression and ileal morphology, including villus structure and goblet cells. In males, menthol partially ameliorated several nicotine-associated changes, including inflammatory markers and barrier-associated genes, consistent with reduced plaque burden, whereas in females, there was evidence of divergent responses. Together, these findings raise the possibility that gut microbial remodeling and altered epithelial barrier function associated with nicotine exposure may contribute to systemic inflammation and increased plaque burden.

## Conclusions

Together, *in vitro* and *in vivo* data suggest females are more sensitive to the vascular effects of vaping across e-liquid formulations, whereas males show partial modulation of select responses, including inflammatory and lipid-associated pathways. Nicotine drove the strongest adverse effects, while menthol attenuated some nicotine-associated effects in a sex-dependent manner but did not eliminate overall vascular or microbial perturbations. Importantly, vehicle exposure alone also altered cardiovascular and microbial outcomes, supporting a role for non-nicotine constituents in e-liquid-associated risk.

A novel finding was that e-liquid exposure altered gut microbial composition and gut barrier-associated pathways in a sex- and treatment-dependent manner, supporting a potential link between microbial remodeling, intestinal dysfunction, and atherosclerosis. The identification of *Alloprevotella* as a discriminating taxon associated with reduced plaque burden further supports a possible gut–vascular axis that warrants future mechanistic investigation.

### Future Directions

Although the ApoE^-/-^ mouse model exhibits elevated basal inflammation that may attenuate treatment effect magnitude, it remains well-suited for evaluating vaping-related atherosclerosis. This model provides a foundation for future studies that can extend vascular phenotyping and further define downstream consequences of e-liquid exposure, including effects on plaque composition and stability, which are critical determinants of cardiovascular events such as myocardial infarction and stroke.

In parallel, the observed shifts in gut microbial communities and gut barrier-associated pathways warrant further investigation to determine whether these alterations contribute causally to e-liquid-associated vascular effects. Future studies incorporating direct measurement of microbial metabolites, including SCFAs and TMAO, as well as microbiota transplantation approaches, may help define the contribution of e-liquids to microbiome remodeling and atherosclerosis progression.

Finally, while the present *in vitro* studies focused on cellular senescence, additional mechanistic analyses examining oxidative stress, autophagy, and vascular cell function would further clarify the molecular pathways underlying the observed responses and their sex-specific differences, building on established links between autophagy, vascular aging, and dysfunction^68^.

